# A statistical note on the detection of autoassociative dynamics in hippocampal place-cell sequences

**DOI:** 10.1101/365239

**Authors:** Lorenz Gönner, Julien Vitay, Fred Hamker

## Abstract

The sequential activity of hippocampal place cells observed during sleep and awake resting is widely viewed as a neural correlate of memory processes. While recent work has advanced our understanding of the content represented during rest-related place-cell sequences, the nature of hippocampal population dynamics during sequential activity remains poorly understood. A recent experimental study has reported that place-cell sequences show a pattern of step-like movement, reminiscent of transitions between discrete attractor states (Pfeiffer and Foster, 2015). By contrast, previous theoretical models predict that the spatiotemporal structure of place-cell sequences should reflect the disribution of place fields, typically observed to be spatially smooth.

Motivated by this discrepancy between models and experimental data, we performed a quantitative comparison between these results and the spike trains generated by a network model for the generation of place-cell sequences (Gönner et al., 2017). Although the model is based on continuous attractor network dynamics, we observed that the movement of sequential place representations was phase-locked to the population oscillation, highly similar to the experimental data interpreted as evidence for discrete attractor dynamics. To resolve this potential contradiction, we performed a detailed analysis of the methodology used to identify discrete attractor dynamics. Our results show that a previous approach to step size decoding is prone to a decoding artefact. We propose a modified approach to estimate step sizes which may help to characterize the underlying circuit dynamics *in vivo*.

## Introduction

The replay of place cells during rest following a spatial experience has been proposed to support both memory recall and memory consolidation, but the underlying mechanisms are largely unknown (Carr et al., 2011). A recent experimental study has sought to determine the population dynamics underlying hippocampal place-cell sequences during sharp wave-ripple events (SWRs) based on large-scale recordings. Pfeiffer and Foster (2015) observed that the location decoded from place-cell sequences moved in an intermittent, step-like fashion, modulated by the phase of slow-gamma (25–50 Hz) population oscillations cooccurring with SWRs: Larger shifts of the decoded location occurred preferentially at times of less population activity. The authors interpreted this effect as evidence for discrete attractor dynamics, in which phases of “sharpening” of a discrete attractor pattern, known as autoassociation, alternate with transitions between different attractor patterns, known as heteroassociation (e.g., Lisman et al., 2005).

This experimental result contrasts with a recent theoretical model for the generation of replay sequences during SWRs, which showed a qualitative pattern of step-like activity transitions, associated with times of low population activity, although the underlying network structure was characerized by a graded pattern of synaptic weights, emerging from sequence learning (Jahnke et al., 2015). This discrepancy between experimental observations and theoretical results is further highlighted by our recent model (Gönner et al., 2017), which shows a qualitative pattern of step-like activity transitions despite continuous attractor dynamics, owing to the presence of strong population oscillations. Moreover, numerous theoretical models for the generation of place-cell sequences predict a spatially smooth structure of sequential activity whenever the spatial distribution of place fields is homogeneous (Redish and Touretzky, 1998, Molter et al., 2007, Hasselmo, 2009, Bush et al., 2010, Vladimirov et al., 2013, Chenkov et al., 2017). By contrast, the generation of step-like sequences would require a spatially discrete clustering of place fields in these models (Bush et al., 2010). To our knowledge, such a clustering has not been observed experimentally (Pfeiffer and Foster, 2013). In brief, while Pfeiffer and Foster (2015) have concluded that discrete attractor dynamics are necessary for generating a spatially discrete pattern of step-like movement, the models of Jahnke et al. (2015) and Gönner et al. (2017) suggest that step-like movement is generated by the interaction between continuously moving sequential representations and strong population oscillations, which impose a temporal discretization on the resulting activity patterns.

To address this apparent discrepancy between theoretical models and recent experimental data, we make a detailed quantitative comparison between simulated spike trains, obtained both using a continuous attractor network model (Gönner et al., 2017) and Poisson activity, and the data reported by Pfeiffer and Foster (2015). Further, to gain a detailed understanding of the decoding method used by Pfeiffer and Foster (2015), we use an analytical approach to characterize its statistical properties. Our results reveal that estimating the step sizes of place-cell sequences represented in oscillatory neural activity is prone to a systematic bias whose magnitude is controlled by the number of spikes per time frame. This decoding artefact arises from the interaction of three factors which would not affect decoding when present individually: (1) Population activity is oscillatory, associated with variable spike counts per decoding frame, (2) cell firing is characterized by random variability (e.g., Poisson-like firing), and (3) step sizes in two-dimensional environments are calculated as the vector length of the *x* and *y* displacements per decoding frame. Under these conditions, which apply to the analysis performed by Pfeiffer and Foster (2015), phase-locked movement (i.e., larger steps at times of low population activity) appears even when the true locations encoded in neural activity change smoothly at a constant speed. These findings suggest that the conclusion of discrete attractor dynamics in the generation of place-cell sequences requires additional substantiation by an analysis which avoids these methodological caveats. We therefore propose a modified approach to step size decoding which strongly reduces the bias of step size estimates for phases of low activity, and which may help to determine whether step-like movement of place-cell sequences is a true effect.

## Results

### An oscillatory continuous attractor network shows phase-locked movement of place-cell sequences

Pfeiffer and Foster (2015) have reported a pattern of step-like movement of hippocampal place-cell sequences at a short timescale, potentially indicative of discrete attractor network dynamics underlying sequence generation. To perform a quantitative comparison between these data and our recent continuous attractor network model for the generation of hippocampal place-cell sequences (Gönner et al., 2017), we reanalysed simulated spike trains generated by our model (see Fig. 1a for an overview of the model architecture). Replicating the analysis described by Pfeiffer and Foster (2015), we performed Bayesian decoding to obtain estimates of represented position during sequential activity, and estimated trajectory step sizes as the vector length of *x* and *y* displacements of decoded location betweeen successive decoding frames.

**Figure 1:**
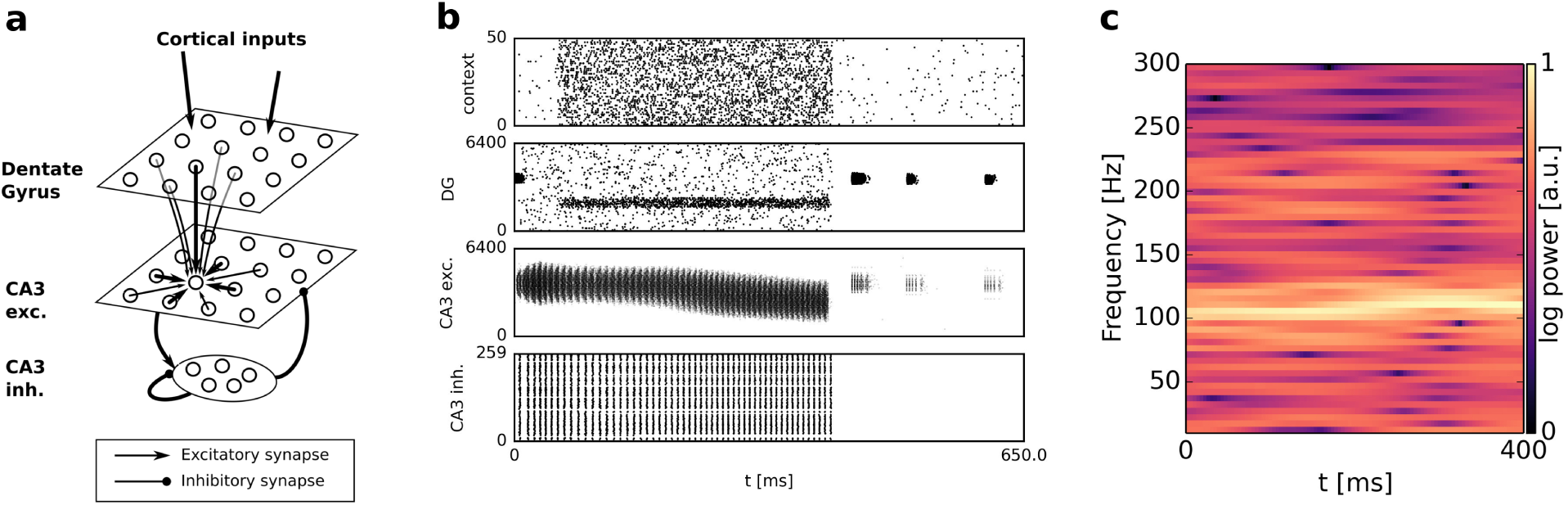
Network architecture and oscillatory dynamics. **a:** Network architecture (simplified). **b:** Example of spiking activity during a simulated place-cell sequence. **c:** Power spectral density of the CA3 population rate during a representative place-cell sequence, indicating oscillatory power in the ripple band. Adapted from Gönner et al. (2017).

In our model, spectral analysis of the population firing rates of CA3 excitatory cells showed strong periodicity in the 100–250 Hz ripple band (Fig. 1c), caused by highly synchronous oscillatory activity in CA3 inhibitory cells (Fig. 1b). We observed that bump movement was phase-locked with the population oscillation in our network (Fig. 2 b, d). Larger movements of the attractor bump and high population activity occurred at opposite phases of the ripple oscillation (Fig. 2 c). In fact, the number of spikes in a given decoding frame was predictive of the size of the bump shift recorded for the same frame (Fig. 2 e).

**Figure 2:**
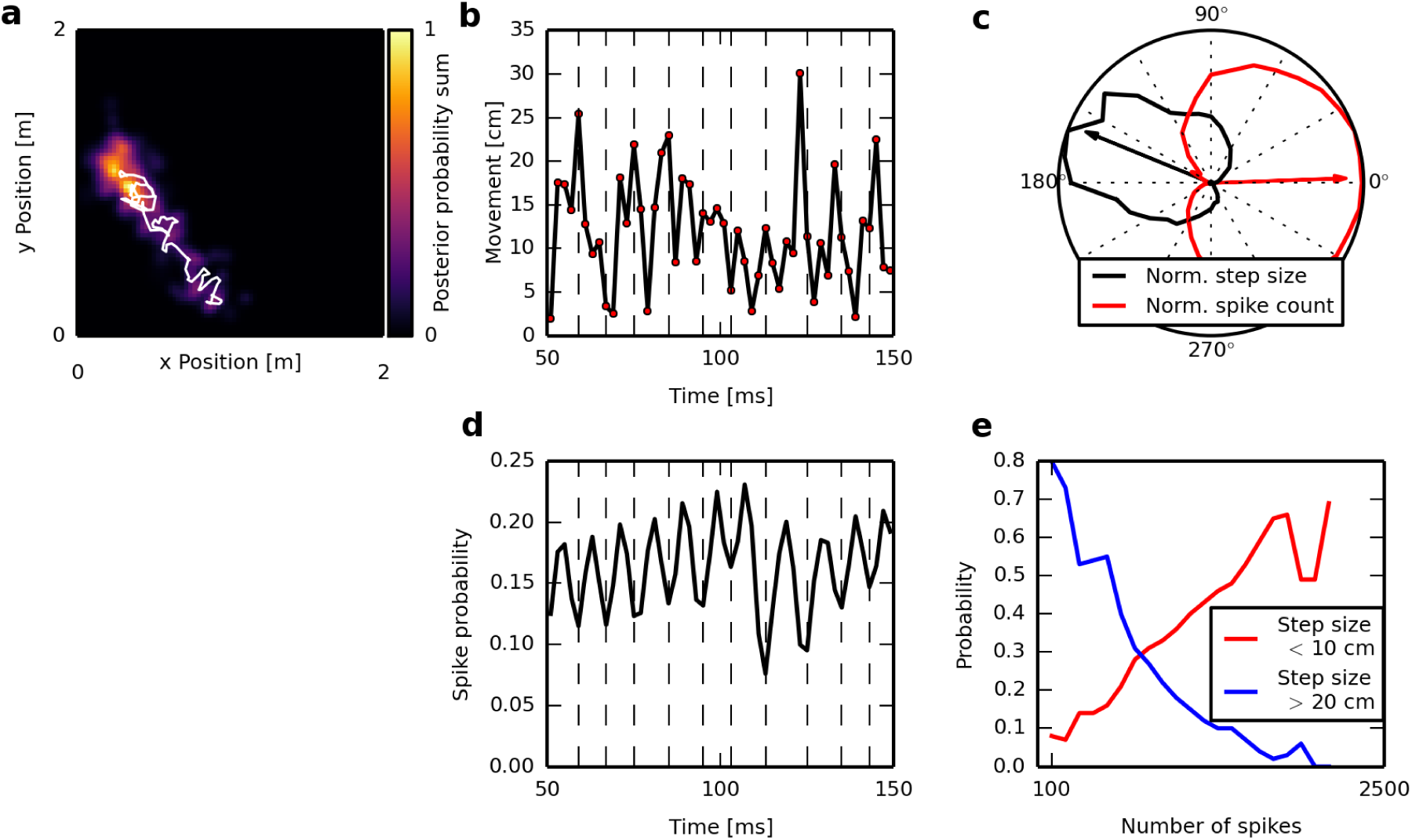
Analysis of bump movement dynamics in an oscillatory continuous attractor network shows a pattern of apparent step-like movement. **a:** Posterior probability sum and trajectory (white line) decoded from a representative sequential event, using a random subset of 160 cells. To improve visibility, decoded positions were averaged across four consecutive decoding frames, totalling 11 ms length. **b:** Bump movement per time frame, defined as the Euclidean distance between the locations decoded from consecutive time frames, for the event shown in (a). **c:** Mean spike count per time frame for the CA3 excitatory population (red) and mean step size per frame (black), as a function of population oscillation phase, normalized to the range [0,1]. Arrows denote the weighted circular mean. **d:** Population activity of CA3 excitatory cells across time, for the event shown in (a). **e:** Probabilities associated with step sizes <10 cm (red) and >20 cm (blue), as a function of population spike count.

Altogether, although we observe ripple-frequency rather than gamma-band oscillations in our model, we have demonstrated that our continuous attractor network shows a substantial modulation of the bump movement speed by the phase of the population oscillation, highly similar to the observations by Pfeiffer and Foster (2015). As this simulation result appears at odds with the conclusion of Pfeiffer and Foster (2015) that phase-locking of bump movement implies discrete attractor dynamics, we investigated the potential origins of this discrepancy.

### Poisson simulations indicate overestimation of step sizes at times of low activity

To further test how accurately the decoded step sizes reflect the true short-term movement dynamics encoded in population activity, we repeated step size decoding on spike trains generated by two control models based on oscillatory Poisson activity. We chose two control models which encoded sequence trajectories with the same overall spatial structure, but which differed in their temporal dynamics of sequence progression: In the “phase-locked movement” model, the speed of ground-truth trajectories was modulated by the phase of the population oscillation; in the “constant movement” model, sequences progressed smoothly at a constant speed (see Methods for details).

In both models, we observed that the accuracy of step size decoding varied considerably across oscillation phases (Fig. 3): While step sizes decoded at times of high population activity provided a good approximation of the true step sizes, we observed that step sizes decoded at times of low activity were substantially larger than their ground-truth values. Consequently, even when the true movement speed was constant across oscillation phases, the magnitude of decoded movement steps appeared to be correlated to the phase of the population oscillation. When plotting individual step size estimates across spike count (more precisely, the harmonic mean spike count across two consecutive time frames; see Methods), we observed a significant negative correlation between step size and spike count, both for true phase-locked movement (*r* = −0.5, *p* < 10^−300^) and for smooth movement (*r* = −0.42, *p* < 10^−300^) (Fig. 4 a,b). However, we found that a 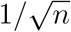 function provided a better fit to these data than a linear fit (*R*^2^ =0.44 vs. 0.25 for phase-locked movement; *R*^2^ =0.31 vs. 0.18 for smooth movement). For comparison, we also investigated the one-dimensional estimates of *x* and *y* displacement. For the model encoding phase-locked movement, we observed a significant negative correlation between decoded *x* and *y* step sizes and mean spike count (*r* = −0.32, *p* < 10^−300^). For smooth movement, we found that *x* and *y* displacements were virtually uncorrelated to mean spike count (*r* = −0.09), consistent with the constant speed of trajectories encoded in this model.

**Figure 3:**
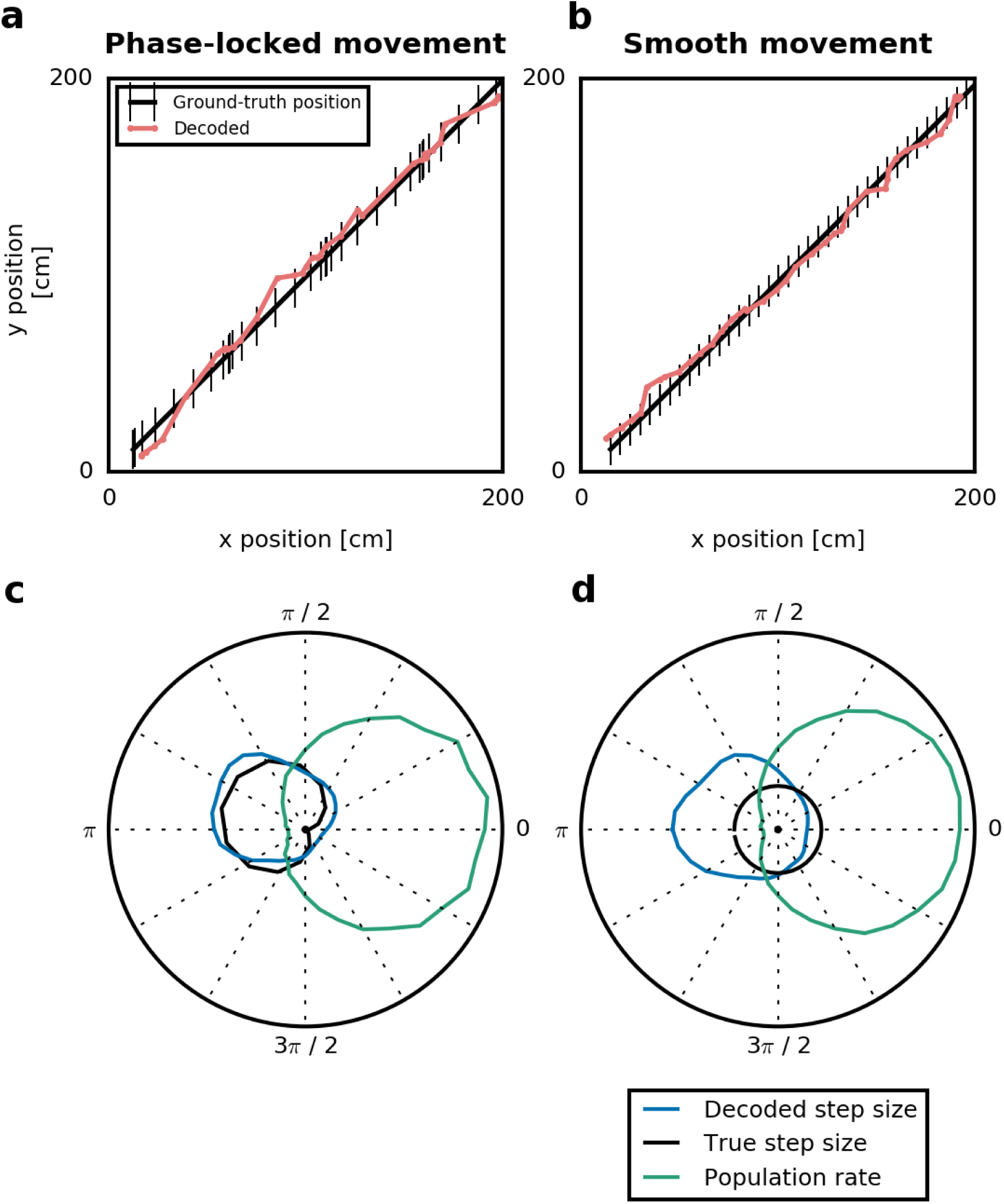
Decoding error of step size estimates varies across oscillation phases. Following a previous approach (Pfeiffer and Foster, 2015), we calculated the step size per time frame as the Euclidean distance (vector length) of the displacement in *x* and *y* directions between successive decoding frames, determined the corresponding phase of the population oscillation, and then averaged the vector lengths within each phase bin, for both phase-locked (a,c) and constant (b,d) movement. **a,b:** Decoded positions versus the true positions encoded by Poisson activity, for representative trajectories. **c,d:** Step size estimates and population activity (as in c and d), averaged averaged across and normalized to a maximum of 1. Note that this approach to estimate mean step sizes shows a high decoding error at times of low population activity, appearing as phase-locked movement even for constant step sizes.

**Figure 4:**
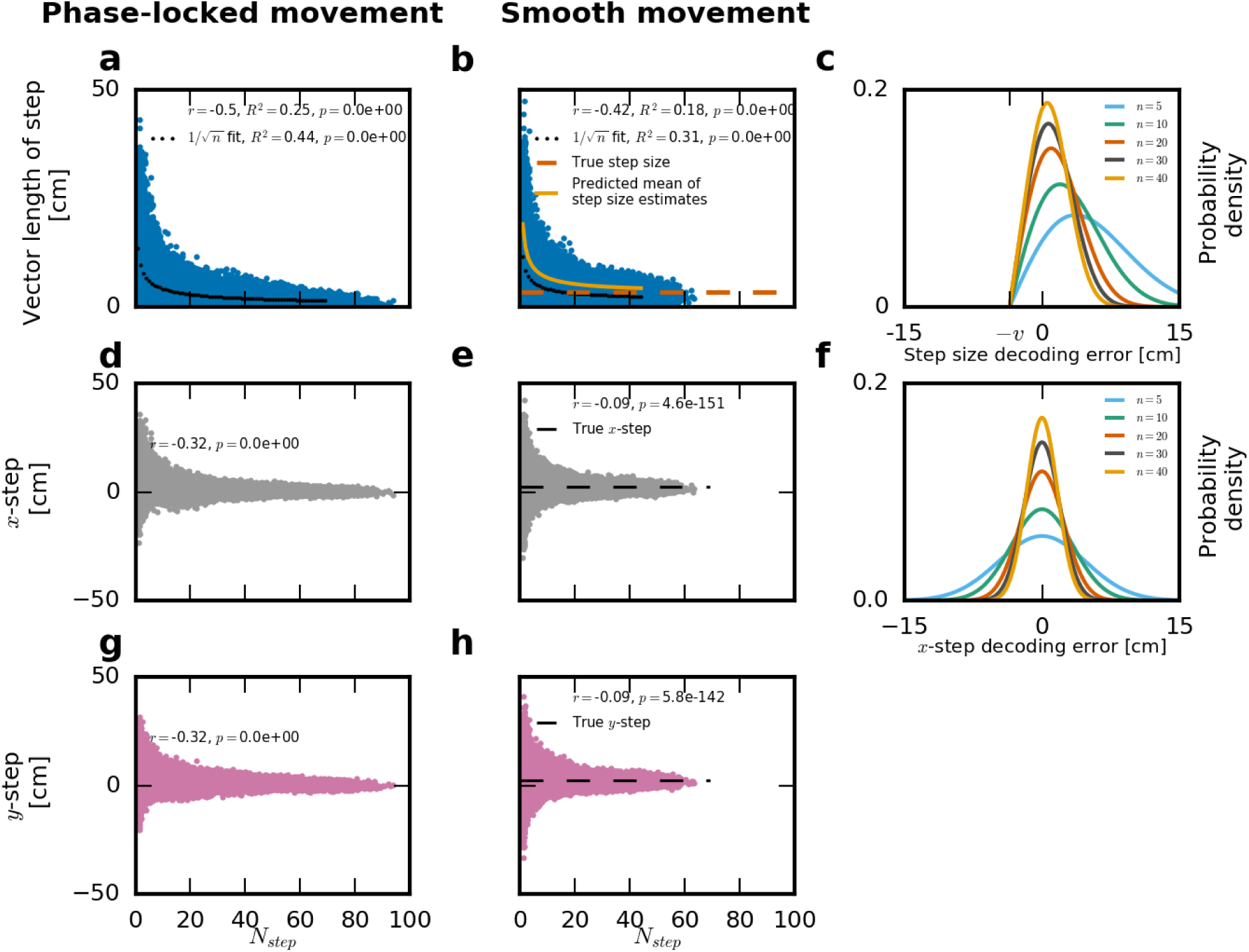
The decoding error of step size estimates is controlled by the number of spikes available for decoding. Simulated data shown for the model encoding phase-locked movement (a,d,g) and for the model encoding smooth movement (b,e,h). **a,b:** Blue dots: Decoded step sizes per time frame, calculated as the vector length of displacement in *x* and *y* directions, plotted across *N_step_* (one-half times the harmonic mean spike count across consecutive decoding frames). Black dots indicate a 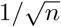 fit to the step size data. Linear correlation coefficients as well as *R*^2^ values for a linear fit and the 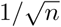 fit shown in the upper part of each figure. In (**b**), the dependence of mean step size estimates on spike count, as predicted by the Rice distribution, is added for comparison (yellow line). **c:** Theoretical distribution of the decoding error of step sizes, in the case of constant movement (based on the Rice distribution). **d,e:** Decoded *x* displacement per time frame, plotted across spike count. **f:** Theoretical distribution of the decoding error of *x*-steps, in the case of constant movement (normal distribution). **g,h:** Decoded *y* displacement per time frame, plotted across spike count. Note that the individual estimates of the *x* and *y* component of step sizes are unbiased (i.e., zero mean decoding error), independent of spike count. By contrast, step size estimates based on the vector length of *x* and *y* steps show a mean decoding error which grows large for low spike counts. Calculated *p*-values of 0.0 indicate a truncation of numerical values below approx. 10^−300^.

Altogether, this indicates a specific limitation of the method of decoding two-dimensional step sizes during times of low population activity, which impairs the discrimination between constant-speed sequence trajectories and phase-locked sequence speeds.

### Biased step size estimates create a confound between decoded step size and spike count

To understand the origins of the reduced accuracy of step size decoding for times of low activity, we took an analytical approach to investigate the statistical properties of step size estimates (see Methods for details). In particular, we derived analytical expressions for the expected distribution of decoded step sizes, assuming sequential place representations progressing at a known speed, encoded in oscillatory Poisson activity.

We first consider the estimated *x* and *y* coordinates of position and step size, which form the basis of two-dimensional step size estimates in an open-field environment. We confirm that the decoded *x* and *y* components of position are unbiased (i.e., their distribution is centered on the true population mean), with a variability which depends on the spike count per decoding frame, *N_spike_*, with a factor of 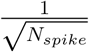. Similarly, the decoded *x* and *y* displacements of position between successive decoding frames are unbiased estimates of the true one-dimensional movement, and their variability is scaled by a factor of 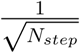, with *N_step_* denoting half the harmonic mean spike count across consecutive decoding frames. However, we observe qualitatively different statistical properties of decoded position and step sizes in the two-dimensional case considered by Pfeiffer and Foster (2015). We note that the decoding error of two-dimensional position estimates, measured as the vector length (Euclidean distance) of the deviation between *x* and *y* position estimates and the respective ground-truth positions, follows a Rayleigh distribution. As such, the mean 2D position decoding error is scaled by a factor of 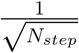, which indicates that 2D position estimates are on average less accurate at times of lower activity. Further, we find that the distribution of 2D step size estimates follows a Rice distribution, a generalization of the Rayleigh distribution. In particular, the mean decoding error of 2D step sizes approaches zero only for large spike counts, while low spike counts are associated with a mean decoding error on the order of 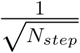. Our simulations confirmed that the Rice distribution provides a good approximation to the mean step size estimates across the range of spike counts (Fig. 4 b).

In summary, we find that estimating step sizes in two-dimensional environments as the vector length of *x* and *y* displacements induces a systematic positive estimation bias controlled by the number of spikes available for decoding. In oscillatory networks, the magnitude of this bias is correlated to oscillation phase, and therefore constitutes a potential confound when comparing decoded step sizes acros oscillation phases.

### Improved step size decoding improves the discrimination between phase-locked and smooth movement

Given that 2D step size estimates used in previous studies are prone to a systematic positive estimation bias, we asked whether step size decoding could be improved to reflect the true movement of represented locations more accurately. As the one-dimensional estimates of *x* and *y* displacements are unbiased, we investigated a modified approach to step size decoding as follows: Instead of averaging across step size estimates based on the vector length of *x* and *y* displacements, we first calculated the mean *x* and *y* displacements for each phase bin, and then took the vector length of these means to estimate the mean step size for this phase bin. We expect this approach to reduce estimation bias, especially for times of low activity, as the number of samples available per phase bin is typically much larger than their mean spike count per decoding frame. Indeed, our simulations confirmed that decoding accuracy increased substantially for the improved estimate, in particular for phases of low activity, although the improved estimates still showed some degree of deviation from the ground-truth step sizes (Fig. 5). Additional analyses indicated that the artefactual correlation between spike count and step size estimates could also be avoided by computing the partial correlation between oscillation phase and step size, controlling for spike count (data not shown). To conclude, we find that artefactual correlations between step size and oscillation phase are reduced when improved step size estimates are considered.

**Figure 5:**
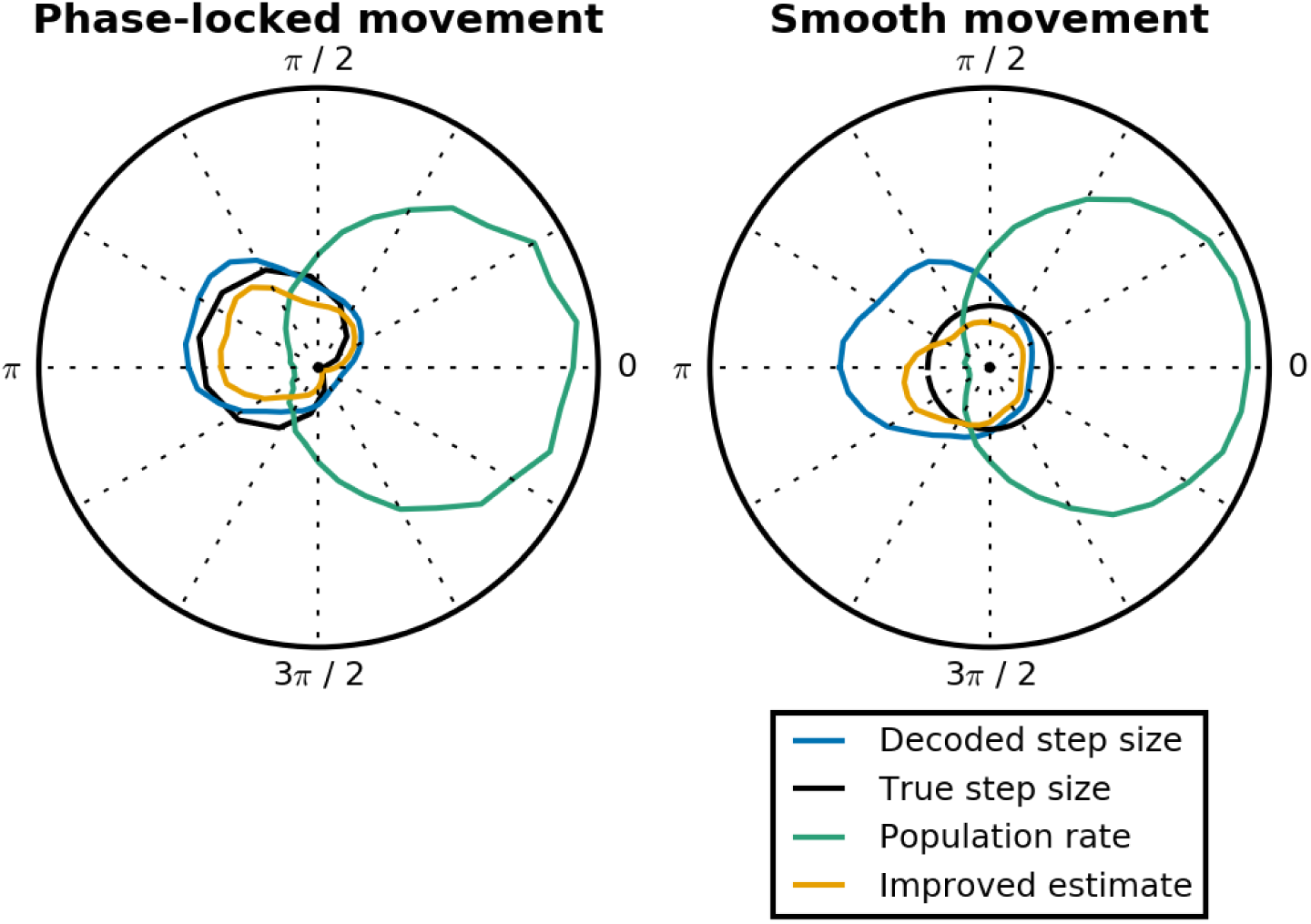
Improved estimates of step sizes discriminate between phase-locked and smooth movement. To calculate an alternative estimate for phase-specific step sizes, we first calculated the arithmetic mean of the displacement in *x* and *y* directions between successive decoding frames for each phase bin. We next determined the mean step size for this phase bin as the Euclidean distance (vector length) of the average *x* and *y* displacements. Data obtained from 1000 simulated sequences with either phase-locked (left) or constant (right) step sizes. Unbiased step size estimates and population activity averaged across oscillation cycles and normalized to a maximum of 1. Note the improved overall agreement between unbiased estimates and true step sizes for the case of smooth movement, which improves the discrimination between phase-locked and smooth movement dynamics.

**Figure 6:**
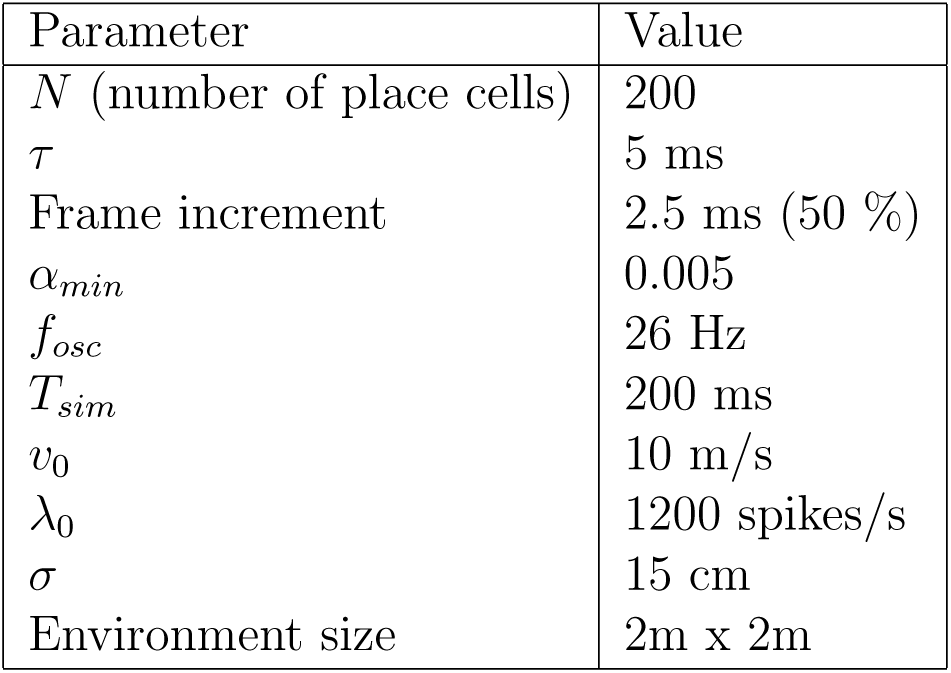
Parameter values for simulations based on Poisson activity.

## Discussion

A recent experimental study obtained large-scale recordings of hippocampal place-cell sequences, allowing to characterize the spatiotemporal patterns of sequential activity at a short timescale. These results showed a pattern of step-like movement of place-cell sequences in awake resting states, consistent with transitions between discrete attractor network patterns (Pfeiffer and Foster, 2015). To perform a quantitative comparison between our recent network model for the generation of place-cell sequences (Gönner et al., 2017) and the experimental data presented by Pfeiffer and Foster (2015), we have repeated their analysis for simulated spike trains. Contrary to our expectations, we observed a high degree of similarity between the results obtained using a continuous attractor network model and the experimental data, interpreted as evidence for discrete attractor dynamics. To resolve this potential contradiction, we performed control analyses based on Poisson simulations, which confirmed that decoded trajectory step sizes differed considerably from ground-truth step sizes. Further, to gain a theoretical understanding of this apparent decoding artefact, we took an analytical statistical approach to investigate the methodology used to estimate trajectory step sizes.

Our main finding is that the choice to estimate step sizes in two-dimensional environments as the vector length of the displacements in *x* and *y* directions is prone to a decoding artefact. This finding holds for network regimes in which spiking activity is stochastic and oscillatory, as is typically the case during recordings of hippocampal sequential activity. Finally, we propose a an alternative approach to step size decoding which provides more reliable estimates of the true step sizes, and which may help to discriminate between discrete and continuous attractor network dynamics in experimental data.

### Limitations

In this theoretical study, we have investigated the analysis methods which may discriminate between discrete attractor dynamics and continuous attractor dynamics in hippocampal place-cell sequences. While theoretical considerations have identified a potential decoding artefact, the magnitude of this artefact is likely scaled by several parameters such as spike rate, the depth of oscillatory modulation of neuronal firing, and decoding window length and overlap. As such, our analysis of the methodological approach used by Pfeiffer and Foster (2015) does not imply that the conclusion based on that method is incorrect. Instead, our results encourage a reanalysis of available experimental data based on an unbiased statistical approach. This way, full advantage may be taken of recent advances in large-scale recording technology to characterize hippocampal circuit dynamics *in vivo*.

## Methods

### Model simulations

The results presented in Fig. 2 have been obtained from spike trains generated by the simulation model of Gönner et al. (2017). As an additional control, we generated spike trains from two models based on oscillatory Poisson activity: A “constant movement” model in which the ground-truth location represented in population activity varied smoothly with constant speed (*v_x_,v*_y_), and a “phase-locked movement” model, in which the progression speed of the ground-truth sequence trajectory was scaled by the phase of the population oscillation.

In both models, we generated population oscillations at a frequency *f_osc_*, by applying a time-varying scaling factor *λ*_peak_(*θ*) to each cell’s firing rate:

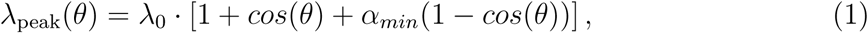

where *λ*_0_ determines the mean firing rate, and *α_min_* controls the modulation depth. For simplicity, we assumed that the variations of *λ*_peak_(*θ*) occurred at a fixed oscillation frequency *f_osc_*:

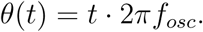

We chose to simulate *N* = 200 place cells, in the range of the cell yield reported by Pfeiffer and Foster (2015). We randomly assigned place field center locations 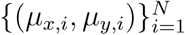 to all cells, based on a uniform distribution in a square environment of size 2m × 2m. We assumed a Gaussian profile of firing rates, with uniform width *σ*.

For a ground-truth location (*x,y*) to be represented in neural activity, the Poisson rates during simulated place-cell sequences were determined by

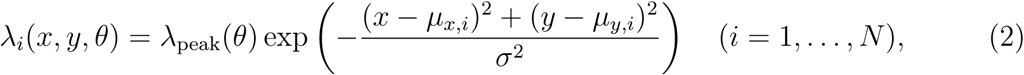

where *λ*_peak_ is the oscillatory scaling factor defined in Eq. (1).

In both models, ground-truth sequence trajectories were defined as

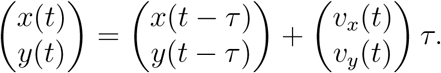

In the “constant movement” model, we set

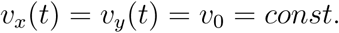

In the “phase-locked movement” model, we assumed sinusoidally varying speeds:

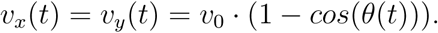

In summary, the generated trajectories were identical in both models, although their progression speed varied.

For both models, we generated 1000 repetitions of independent Poisson spike trains defined by the ground-truth sequence trajectories. For an overview of parameter values used in simulations, see Table 6.

### Statistical properties of Bayesian location estimates

The posterior probability of the location *X* represented in neural activity to be a potential location *x* out of a set of position bins 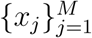, given an observation 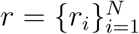 of neural activity *R*, is

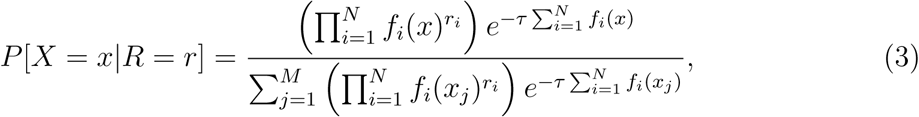

where *f_i_* is the spatial tuning curve of unit *i* (in our case, a Gaussian function with width *σ*), *r_i_* is its spike count, and *τ* is the length of the decoding window. This approach assumes that all *N* units follow independent Poisson firing statistics, and that occupancy is uniform across locations (Davidson et al., 2009).

It has been shown that the uniform-prior Bayesian estimate is equal to the maximum-likelihood place estimate *x_est_* (Dayan and Abbott, 2001). In the present case of Gaussian-shaped place fields, this estimate has the convenient structure

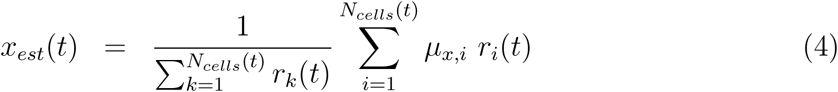

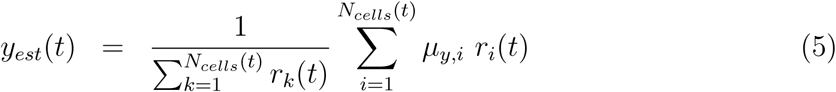

(Dayan and Abbott, 2001).

The distribution of *x_est_*(*t*) is therefore a weighted sum of Poisson random variables, where both the Poisson rates and the weight coefficients are determined by the Gaussian function describing the population code. Although we are not aware of a closed-form expression of this type of distribution, we note that Eqs.(4) and (5) can be rewritten in a simplified form as

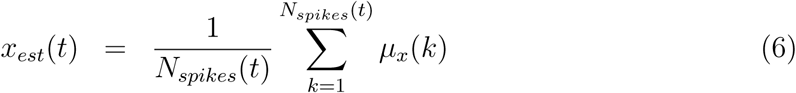

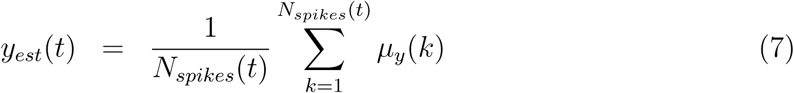

where (*μ_x_*(*k*),*μ_y_*(*k*)) denotes the place field center location of the cell associated with spike *k* ∈ {1, …, *N_spikes_* (*t*)}, *r_i_* is the spike count of cell *i*, and 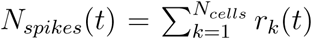. This motivates the following approximation to characterize *x_est_*(*t*): By our assumptions on the population code, the place field center locations *μ_x_* associated to the recorded spikes can be viewed as samples from a normal distribution with mean *x*(*t*) and variance *σ*^2^. Eq. (6) reveals that *x_est_*(*t*) follows a sampling distribution, which in this case is a normal distribution, with mean:

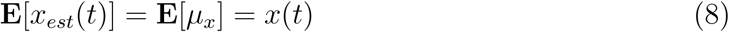

and variance:

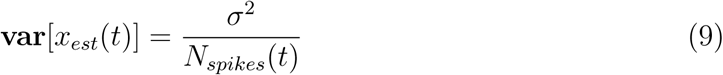

(Shao, 2003), where **E**[.] denotes expectation, and **var**[.] denotes variance. The same properties hold for *y_est_*(*t*) in analogy. In other words, the one-dimensional position estimates *x_est_* and *y_est_* are unbiased, with a variance which depends on the number of spikes available for decoding.

### Statistical properties of step size estimates

Throughout this section, we assume that decoding windows are non-overlapping, to ensure that consecutive place estimates *x_est_*(*t*) and *x_est_*(*t* − *τ*) are statistically independent.

We define the estimates of *x*-step sizes as

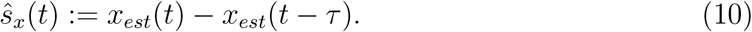

Due to the additive property of the variance,

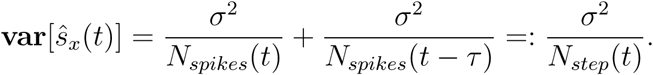

As *ŝ_x_*(*t*) is a weighted sum of Gaussian random variables, *ŝ_x_*(*t*) follows a normal distribution with mean *v_x_τ* and variance 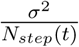, where *v_x_τ* is the true *x*-displacement per decoding frame, and

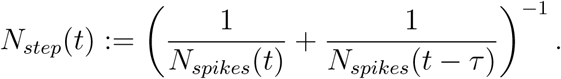

(Note that *N_step_*(*t*) is equal to one-half times the harmonic mean of *N_spikes_*(*t*) and *N_spikes_*(*t* − *τ*).) The estimate of *y*-step sizes, *ŝ_y_*(*t*), is defined in analogy, with *v_y_τ* denoting the true *y*-displacement per decoding frame. The decoding error of *x*-step estimates, denoted by Δ_*Sx*_, equals the difference between the *x*-step estimate and the true *x*-step:

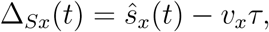

and, defining Δ_*Sy*_ analogously as the decoding error of the *y* step,

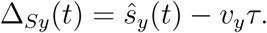

Hence, Δ_Sx_(*t*) and Δ_Sx_(*t*) are both normally distributed, with mean zero and variance 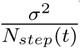. As an intermediate result, it follows that the vector length of these decoding errors of *x* and *y* steps follows a Rayleigh distribution with parameter 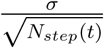 (see section A.0.1 and Fig. 7). In particular, the mean decoding error equals 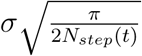, i.e., is proportional to 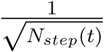.

**Figure 7:**
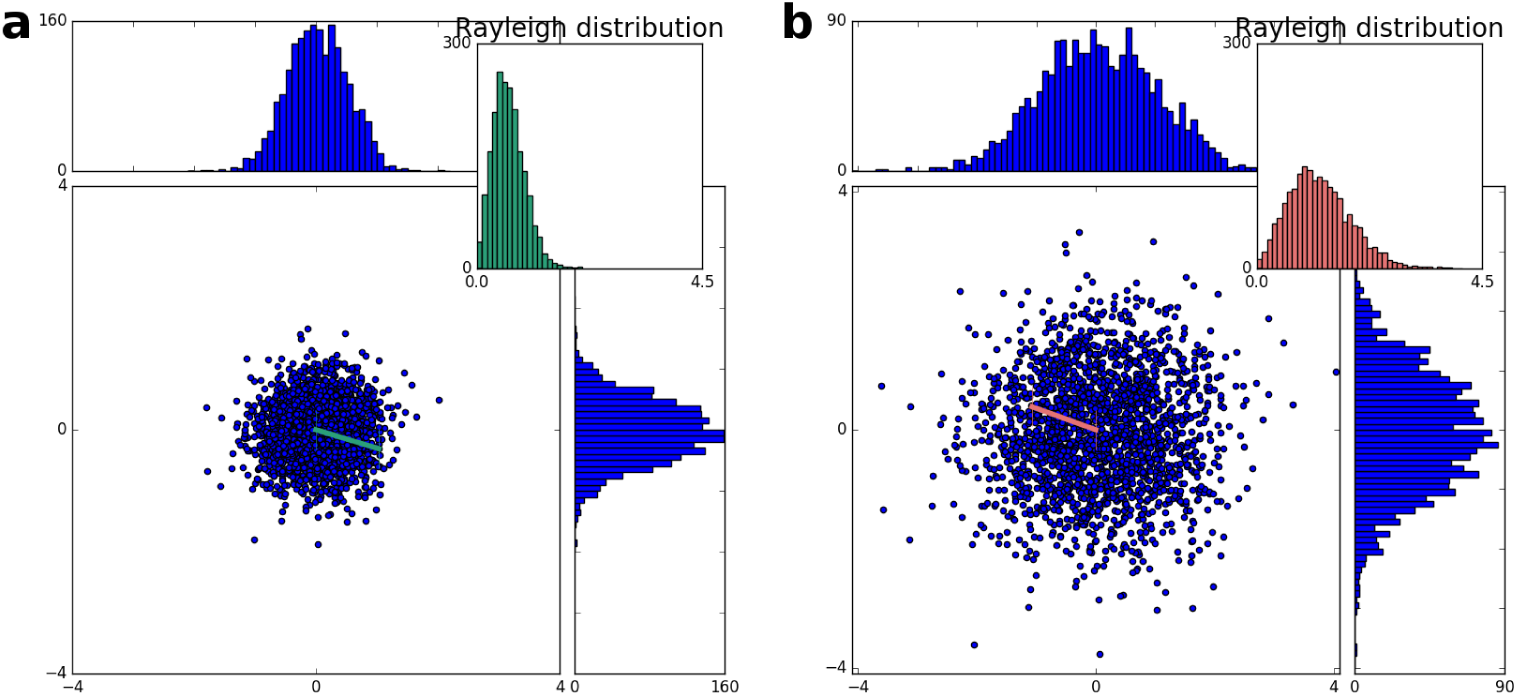
Illustration of the Rayleigh distribution. The Rayleigh distribution describes the vector length (Euclidean distance) of samples from a two-dimensional Gaussian random variable with zero mean and identical standard deviation *σ*. **a:** *σ* = 0.5, **b:** *σ* = 1. Note the increased mean of the Rayleigh distribution in b, caused by larger variability of the Gaussian components.

We next turn to characterize the two-dimensional step size estimates used to analyse the movement dynamics of place-cell sequences. We define the joint (*x,y*)-step size estimate *ŝ_xy_* as the vector length of *x*- and *y*-displacements:

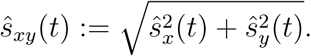

We note that the step size estimate *ŝ_xy_*(*t*) follows a Rice distribution with parameters 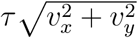 and 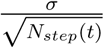, which has a mean of

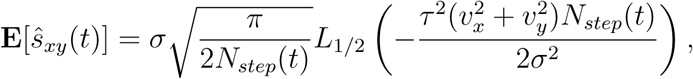

where *L*_1/2_ denotes the Laguerre polynomial (see section A.0.2). Importantly,

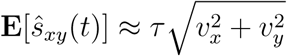

in the limit of infinitely large *N_step_*(*t*) (i.e., the estimator converges to the true step size asymptotically). By contrast, for small values of *N_step_*(*t*), the Laguerre polynomial approximates identity, and therefore

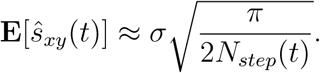

In summary, the two-dimensional step size estimate *ŝ_xy_* is unbiased only in the limit of large spike counts, whereas lower spike counts are associated with a positive estimation bias proportional to 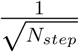. By definition, *ŝ_xy_* is therefore a biased estimator of the true two-dimensional step size 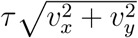.

## A. Supplemental Information

### A.0.1. Rayleigh distribution

If *X* and *Y* are independent normally distributed random variables with mean zero and variance *σ*^2^, then 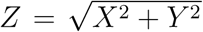 follows a Rayleigh distribution with parameter *σ*. Its probability density function (pdf) is given by

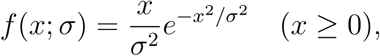

and its mean equals 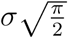 (Siddiqui, 1962).

### A.0.2. Rice distribution

The Rice distribution is a generalization of the Rayleigh distribution for non-zero mean components. It was originally developed as a model for the propagation of noisy radio signals (Rice, 1945).

If *X* and *Y* are independent normally distributed random variables with mean values *ν_x_* and *ν_y_*, then 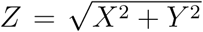 follows a Rice distribution with parameters 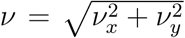 and *σ*.

The pdf of the Rice distribution with parameters *ν* and *σ* is given by

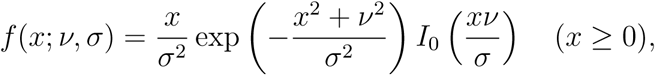

where *I*_0_ denotes the modified Bessel function of the first kind with order zero (Rice, 1945, p.112). Its mean is given by 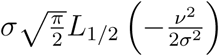. Here, *L*_1/2_ denotes the Laguerre polynomial, which can be expressed as

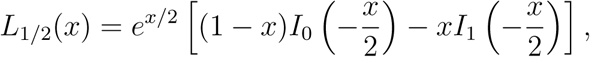

where *I*_1_ denotes the modified Bessel function of the first kind with order one (Rice, 1945, p.113).

We note that in the case of small *σ* or large *v*, the mean value of the Rice distribution approaches *v*, as 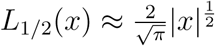 in the limit of *x* → −∞ (Abramowitz and Stegun, 1972, § 13.5.1).

For *v* = 0, the Rice distribution equals the Rayleigh distribution, as *I*_0_(*x*) = 1 for *x* → 0 and *L*_1/2_(*x*) = 1 for *x* → 0 (Abramowitz and Stegun, 1972, § 9.6.7, § 13.5.5).

## References

Abramowitz, M. and Stegun, I. A. (1972). Handbook of mathematical functions. 10th printing, National Bureau of Standards.

Bush, D., Philippides, A., Husbands, P., and O’Shea, M. (2010). Dual coding with STDP in a spiking recurrent neural network model of the hippocampus. PLoS Comput Biol, 6(7):e1000839.

Carr, M., Jadhav, S., and Frank, L. (2011). Hippocampal replay in the awake state: a potential substrate for memory consolidation and retrieval. Nat Neurosci, 14:147–153.

Chenkov, N., Sprekeler, H., and Kempter, R. (2017). Memory replay in balanced recurrent networks. PLoS Comput Biol, 13(1):e1005359.

Davidson, T., Kloosterman, F., and Wilson, M. (2009). Hippocampal replay of extended experience. Neuron, 63:497–507.

Dayan, P. and Abbott, L. (2001). Theoretical Neuroscience. MIT Press.

Gönner, L., Vitay, J., and Hamker, F. (2017). Predictive place-cell sequences for goal-finding emerge from goal memory and the cognitive map: A computational model. Front Comput Neurosci, 11:84.

Hasselmo, M. (2009). A model of episodic memory: Mental time travel along encoded trajectories using grid cells. Neurobiol Learn Mem, 92:559–573.

Jahnke, S., Timme, M., and Memmesheimer, R.-M. (2015). A unified dynamic model for learning, replay, and sharp-wave/ripples. J Neurosci, 35:16236–16258.

Lisman, J., Talamini, L., and Raffone, A. (2005). Recall of memory sequences by interaction of the dentate and CA3: A revised model of the phase precession. Neural Networks, 18:1191–1201.

Molter, C., Sato, N., and Yamaguchi, Y. (2007). Reactivation of behavioral activity during sharp waves: A computational model for two stage hippocampal dynamics. Hippocampus, 17:201–209.

Pfeiffer, B. and Foster, D. (2013). Hippocampal place-cell sequences depict future paths to remembered goals. Nature, 497:74–79.

Pfeiffer, B. and Foster, D. (2015). Autoassociative dynamics in the generation of sequences of hippocampal place cells. Science, 349:180–183.

Redish, A. and Touretzky, D. (1998). The Role of the Hippocampus in Solving the Morris Water Maze. Neural Computation, 10:73–111.

Rice, S. (1945). Mathematical Analysis of Random Noise Part III: Statistical properties of random noise currents. Bell System Technical Journal, 23:46–156.

Shao, J. (2003). Mathematical statistics. 2nd ed. Springer, New York.

Siddiqui, M. (1962). Some problems connected with Rayleigh distributions. Journal of Research of the National Bureau of Standards, Section D: Radio Propagation, 66D:167.

Vladimirov, N., Tu, Y., and Traub, R. (2013). Synaptic gating at axonal branches, and sharp-wave ripples with replay: a simulation study. Eur J Neurosci, 38:3435–3447.

